# Sustained Activation of PV+ Interneurons in Core Auditory Cortex Enables Robust Divisive Gain Control for Complex and Naturalistic Stimuli

**DOI:** 10.1101/832642

**Authors:** Tina Gothner, Pedro J. Gonçalves, Maneesh Sahani, Jennifer F. Linden, K. Jannis Hildebrandt

## Abstract

Sensory cortices must flexibly adapt their operations to internal states and external requirements. Modulation of specific inhibitory interneurons may provide a network-level mechanism for adjustments on behaviourally relevant timescales. Understanding of the computational roles of such modulation has mostly been restricted to phasic optogenetic activation and short, transient stimuli. Here, we aimed to extend the understanding of modulation of cortical inhibition by using sustained, network-wide optogenetic activation of parvalbumin-positive interneurons in core auditory cortex to study modulation of responses to transient, sustained, and naturalistic stimuli. We found highly conserved spectral and temporal tuning, despite profoundly reduced overall network activity. This reduction was predominantly divisive, and consistent across simple, complex, and naturalistic stimuli. A recurrent network model with power-law input-output functions replicated our results. We conclude that modulation of parvalbumin-positive interneurons on timescales typical of more sustained neuromodulation may provide a means for robust divisive gain control conserving stimulus representations.

## Introduction

Sensory processing in the cortex requires flexible and reliable mechanisms for adjusting computations and information flow according to context. Both internal states and external requirements can reliably trigger changes of the most fundamental cortical computations, and these modulations can also generalise across stimulus conditions. The different subtypes of cortical inhibitory interneurons are candidate mediators of contextual modulation because they are in an exquisite position to produce such changes (Markram et al. 2004; Kepecs and Fishell 2014; Tremblay et al. 2016; Cardin 2019).

Cortical interneurons can be subdivided into three classes: parvalbumin positive (PV+), somatostatin positive (SOM+), and those expressing vaso-intestinal protein (VIP+). Each of these classes is differentially targeted by neuromodulation (Swanson and Maffei 2019), and has its unique, class-specific expression profile of neuromodulatory receptors (Paul et al. 2017), resulting in class-specific modulation of interneurons in specific contexts and behavioral contexts (Kawaguchi and Shindou 1998; Toussay et al. 2013; Polack et al. 2013; Poorthuis et al. 2014; Alitto and Dan 2012; Pakan et al. 2016). In order to obtain a functional understanding of neuromodulation, a more detailed description of network effects of the activation of specific interneuron classes is desirable (Edeline 2012). Great progress on our understanding of functional role of modulation of PV+ cell and other classes has been made with the help of optogenetic tools recently (Wilson et al. 2012; Lee et al. 2012; Hamilton et al. 2013; Aizenberg et al. 2015; Seybold et al. 2015; Moore et al. 2018; Atallah et al. 2012), indicating that PV+ may play an important, albeit complex role in controlling neural excitability in sensory cortices.

So far, most studies using precise class-specific manipulation of inhibitory populations have focused on fast time scales, typically locked to sparse and transient stimuli. However, contextual modulation may act at either fast sub-second time scales (McGinley et al. 2015; Muñoz and Rudy 2014; Poorthuis et al. 2014) or over the time course of several seconds to minutes (Kawaguchi and Shindou 1998; Metherate et al. 1992; Alitto and Dan 2012; Schneider et al. 2014; Petrie et al. 2005), and there is evidence that PV+ inhibition may mediate very different network dynamics in sensory cortices, largely depending on the time scales of stimulation (Ozeki et al. 2009; Haider et al. 2010). Thus, in order to investigate the functional role of modulation of PV+ activity at all relevant time scales, a precise and more sustained manipulation of PV+ cell activity is needed in addition to the already available fast and transient optogenetic activation or deactivation (Wilson et al. 2012; Lee et al. 2012; Hamilton et al. 2013; Aizenberg et al. 2015; Seybold et al. 2015; Moore et al. 2018; Atallah et al. 2012). Here, we employed a variant of Channelrhodopsin 2 that allows for sustained, low-level activation at the time scale of minutes (stable-step function opsin, SSFO, (Berndt et al. 2009; Yizhar et al. 2011) to ask how the sustained activation of PV+ cells, mediates control of cortical computation.

We hypothesised that sustained, low-level activation of PV+ cells (Tremblay et al. 2016) provides a means for divisive scaling of neural responses in core auditory cortex. Divisive scaling has been proposed to be one of the canonical cortical computations (Carandini and Heeger 2011), its action spanning from context-dependent processing of sensory stimuli (Rabinowitz et al. 2011; Rabinowitz et al. 2013) to task-dependent top-down control (Ruff and Cohen 2017). Concretely, in order to serve as a general instrument for divisive scaling across a cortical area such a mechanism should (1) provide divisive changes across single-unit responses, (2) leave basic response properties such as tuning and receptive field-structure intact, and (3) generalise across different stimulation paradigms, including complex and naturalistic stimulus sets.

In order to investigate a potential modulatory role of PV+ cells, we expressed SSFO in PV+ cells in the core auditory cortex (AC) of mice. We recorded responses of populations of single units to three different stimulus paradigms: single tones (ST), dynamic random chords (DRC) and a set of animal vocalizations (natural stimuli, NS) in awake mice. The combination of sustained PV+ cell activation and reliable long-term recording allowed us to test whether single units are altered divisively across the different paradigms and to quantify their stimulus encoding properties.

We show that sustained activation of PV+ cells in auditory cortex provides a means for coherent divisive scaling, and changes tuning properties only marginally. This divisive scaling generalises across complex and naturalistic stimuli on both population and single-unit levels. A network model with power-law input-output functions captures these key experimental findings. Overall, our findings provide evidence that sustained activation of PV+ interneurons may constitute a powerful neural instrument for context-dependent scaling of cortical responses.

## Results

### Consistent effects of sustained activation of PV+ cells across trials and paradigms

Many behavioural and brain-state dependent modulations of PV+ activity occur over timescales of several seconds to minutes (Kawaguchi and Shindou 1998; Metherate et al. 1992; Alitto and Dan 2012; Schneider et al. 2014; Petrie et al. 2005; Castro-Alamancos and Connors 1996). In order to mimic such modulations and to probe their functional significance for cortical computation at the network level, we decided to employ a stable step-function variant of ChR2 (stable step-function opsin, SSFO) expressed in PV+ interneurons. SSFO allows for constant depolarisation of the targeted cells for minutes after applying a single, short pulse of light (Berndt et al. 2009; Yizhar et al. 2011). This mode of action not only allowed for more modulatory-like activation of PV+ cells, but enabled us to examine the effect of PV+ cell activation on the encoding of extended auditory stimuli of greater complexity. Thus, we could compare effects of PV+ activation during prolonged dynamic random chord (DRC) stimuli to effects observed with more traditional single-tone (ST) paradigms (Seybold et al. 2015; Aizenberg et al. 2015; Phillips and Hasenstaub 2016; Moore et al. 2018).

To determine whether sustained activation of PV+ interneurons produces consistent changes in neural responses across trials and stimulus paradigms, we first compared overall firing rates of single-unit responses to ST and DRC stimuli recorded in awake mice (Fig. 1A) with (SSFO•PV) and without (control) activation of PV+ cells. Using blue and orange light to activate/deactivate SSFO, we found a robust and reproducible effect of sustained PV+ activation in single units (Fig. 1B). Among the recorded units, we observed both robust decreases and increases in firing rate (Fig. 1C). These changes were consistent for ST and DRC stimuli when looking at overall firing rates during either stimulus paradigm (correlation coefficient r = 0.42, p = 2e-63, n = 1422) with no systematic difference in single-unit modulation under DRC and ST stimulation (Fig. 1D, p = 0.367, Wilcoxon rank-sum test, n = 1422). At the population level, we observed on average a decrease in population firing rate during PV+ activation for both ST (mean (SD) rate change = −22.93% (±67.79%)) and DRC stimulation (mean (SD) rate change = −26.98% (±36.18%)). Effects of sustained PV+ activation on firing rates in single units ranged continuously from complete suppression to robust elevation.

**Figure 1.**
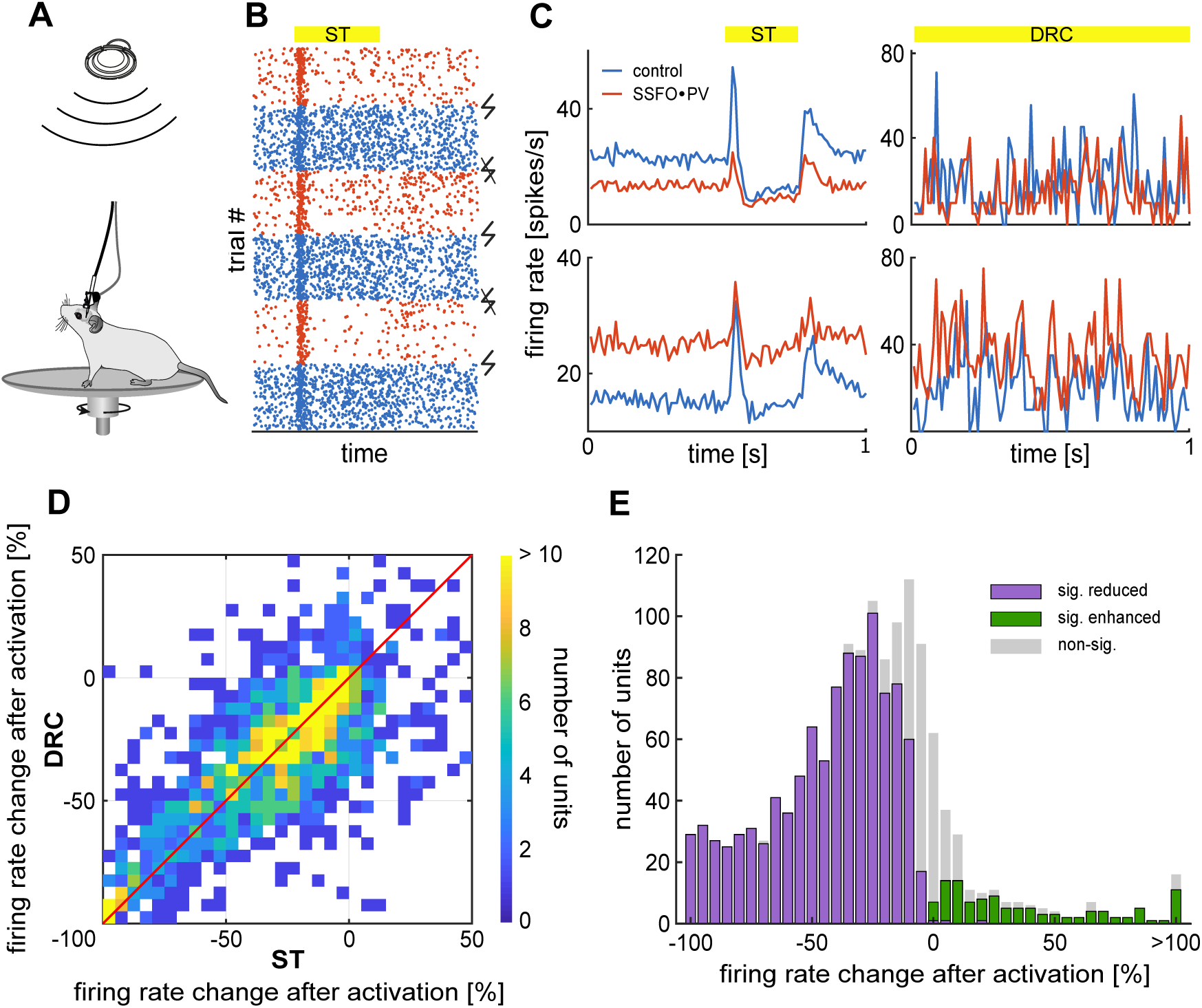
Sustained activation of PV+ cells results in consistent changes of firing rates across trials and paradigms. (**A**): Recording setup. Mice were implanted with a combined opto-electrode implant and were able to freely move on a horizontal running wheel while sound was presented from above. (**B**): Responses of an example unit to sound (yellow bar) in control (blue) and SSFO•PV trials (red). Spiking activity changes immediately after activation of PV+ cells (flash) and lasts until deactivation (strikeout flash). (**C**): Example PSTHs of two units (top, bottom) during single tone stimuli (ST, left) and dynamic random chords (DRC, right). PV+ activation resulted both in decreased (top) and increased (bottom) firing rates in individual units. (**D**): Comparison of the percentage firing rate change in individual units after PV+ activation during ST and DRC (n = 1422 units).The red diagonal line indicates equal firing rate changes with PV+ activation in both stimulus paradigms. (**E**): Distribution of percentage firing rate change after PV+ activation during ST stimulation. Classification of units into two groups for further analysis: units with a significantly reduced firing rate (reduced units, purple, n = 1027) and units with a significantly enhanced firing rate (enhanced units, green, n = 111) during sustained PV+ activation.

### Are the enhanced units PV+ interneurons?

A small proportion of the recorded units showed an enhancement of firing rate with sustained PV+ activation rather than a reduction (Fig. 1). Because SSFO depolarizes directly activated cells, we hypothesized that a large proportion of the enhanced cells were directly-activated PV+ cells rather than pyramidal cells or non-PV+ inhibitory neurons in the cortical network. In order to test this, we analyzed extracellular spike shapes separately for significantly enhanced and reduced units.

Previous work reported that most of PV+ cells have narrow spike duration (fast-spiking cells, Keller et al. 2018) and an increased peak-to-trough ratio (Kim et al. 2016; Moore and Wehr 2013), criteria that we used to find putative PV+ cells in the group of enhanced units. We extracted spike width and trough amplitude in normalized waveforms and compared these between reduced and enhanced units (Fig. 2 A). Mean trough amplitude for the enhanced units was significantly larger (Fig. 2 B, top; mean (SD) = 0.2513 (± 0.2285)) than for reduced units (mean (SD) = 0.1522 (± 0.1713), Tukey-Kramer Test, p = 0.0021). Furthermore, enhanced units did have significantly smaller mean spike widths (Fig. 2 B, bottom; mean (SD) = 0.1804 ms (± 0.0552 ms)) compared to reduced units (mean (SD) = 0.2190 ms (± 0.0741 ms), Tukey-Kramer Test, p = 1.16e-4). The distribution of spike widths for enhanced units exhibited a local minimum at approximately 0.12 ms. We observed substantially different optogenetic modulation for neurons with spike widths falling above and below this value (Fig. 2 C). Units with a spike width larger than 0.12 ms exhibited a highly significant reduction in firing rate during sustained PV+ activation (mean (SD) = −40.64 % (± 38.88 % ms), Mann-Whitney-U-Test, p = 3.14e-15) compared to units with a smaller spike width (mean (SD) = −5.61 % (± 32.82 % ms)). Thus, we found clear evidence for a relationship between spike shape and rate modulation during SSFO•PV trials. However, we did not find a clear bimodal distribution of any of these features or a combination of them in our data (Fig. 2 D). Narrow spiking units were more often enhanced than other units, but often reduced in their firing. Similar mixed effects on PV+ cells due to indirect network effects have been reported previously (Moore and Wehr 2013).

**Figure 2.**
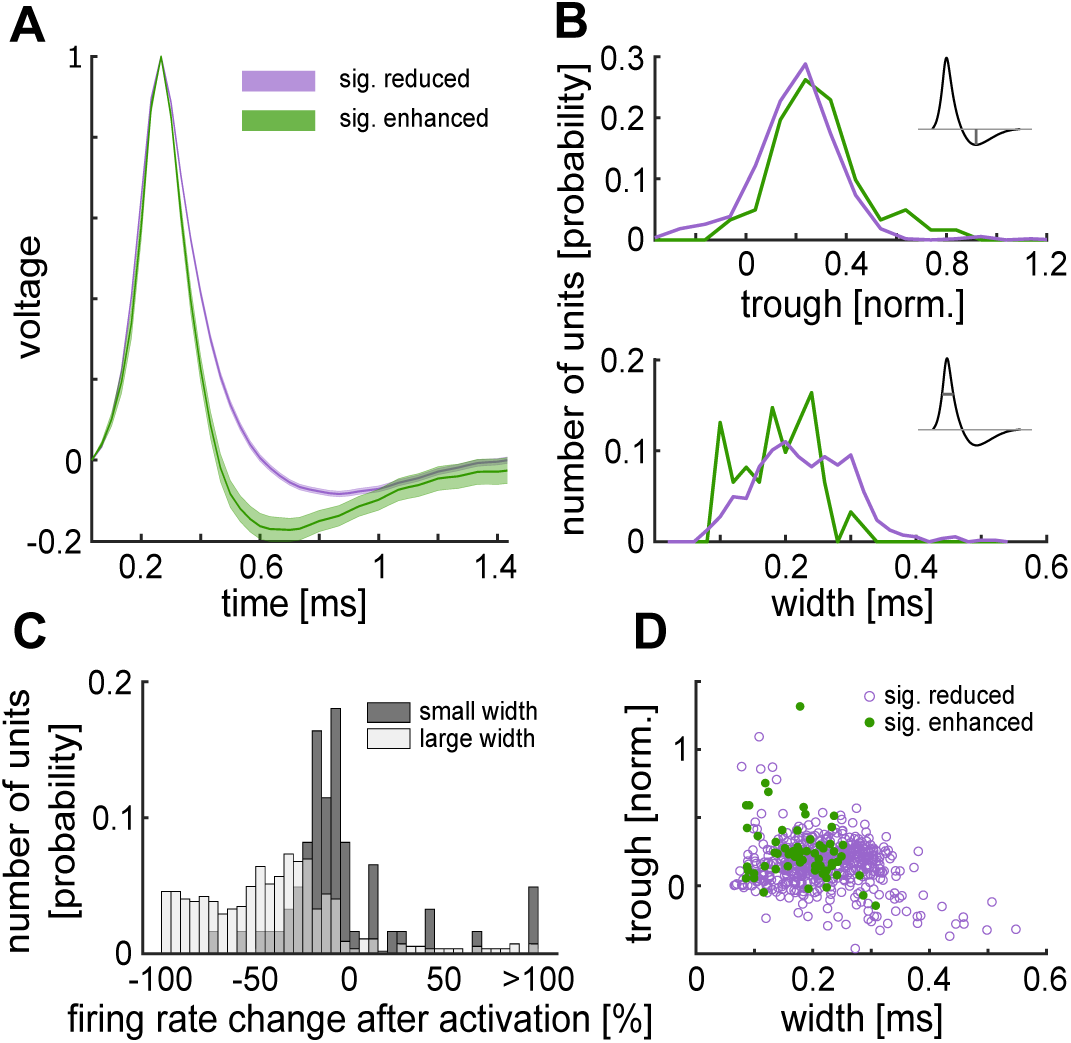
Reduced and enhanced units display different spike shapes. (**A**): Mean normalized waveforms grouped into reduced (purple, n =45) and enhanced (green, n = 61) units. Shaded area depicts SEM. (**B**): Histograms of trough amplitude (top, normalized to peak) and spike width (bottom, at 50% of peak amplitude) within the subgroups. (**C**): Distribution of firing rate changes after PV+ activation for units with a spike width smaller (light grey, n = 61) and larger (dark grey, n = 545) than 0.12 ms. (**D**): Relationship of width and trough for reduced (purple, n = 545) and for enhanced units (green, n = 61).

Since no clear separation between putative PV+ and PV-units was possible, we decided to group units on a functional basis reflecting the direction of firing rate change instead (reduced units; n = 1027; enhanced units; n = 111, Fig. 1E). Based on the analysis of spike shapes, we expect the enhanced group to contain mostly directly activated PV+ cells, but many PV+ cells to be contained in the reduced group as well.

### Activation of PV+ cells preserves frequency tuning properties of neurons in auditory cortex

We next asked whether sustained PV+ activation would conserve basic encoding properties in auditory cortex, as would be expected for general divisive gain control mechanisms harnessed by contextual modulation, or whether it might instead sharpen or shift tuning. To address this question we recorded responses to isolated single tones of varying frequency. We constructed tuning curves using the onset responses of the units for both control and SSFO•PV conditions.

We observed a wide range of changes to the tuning curves in the recorded units (Fig. 3). Fig. 3A depicts the change in tuning curves after PV+ activation, ordered by the change at best frequency (BF). In the group with overall enhanced units, units could show increased firing rates over all test frequencies (example in the lower left box, Fig. 3A) or even a reduction of firing at BF but increased rate at off-BF sound frequencies (Fig. 3A, upper left example). A similar range of effects could be observed in the group with overall reduced firing rates: some units showed a general reduction of firing rates (Fig. 3A, upper right example), others were reduced mostly at off-BF frequencies (Fig. 3A, lower right example).

**Figure 3.**
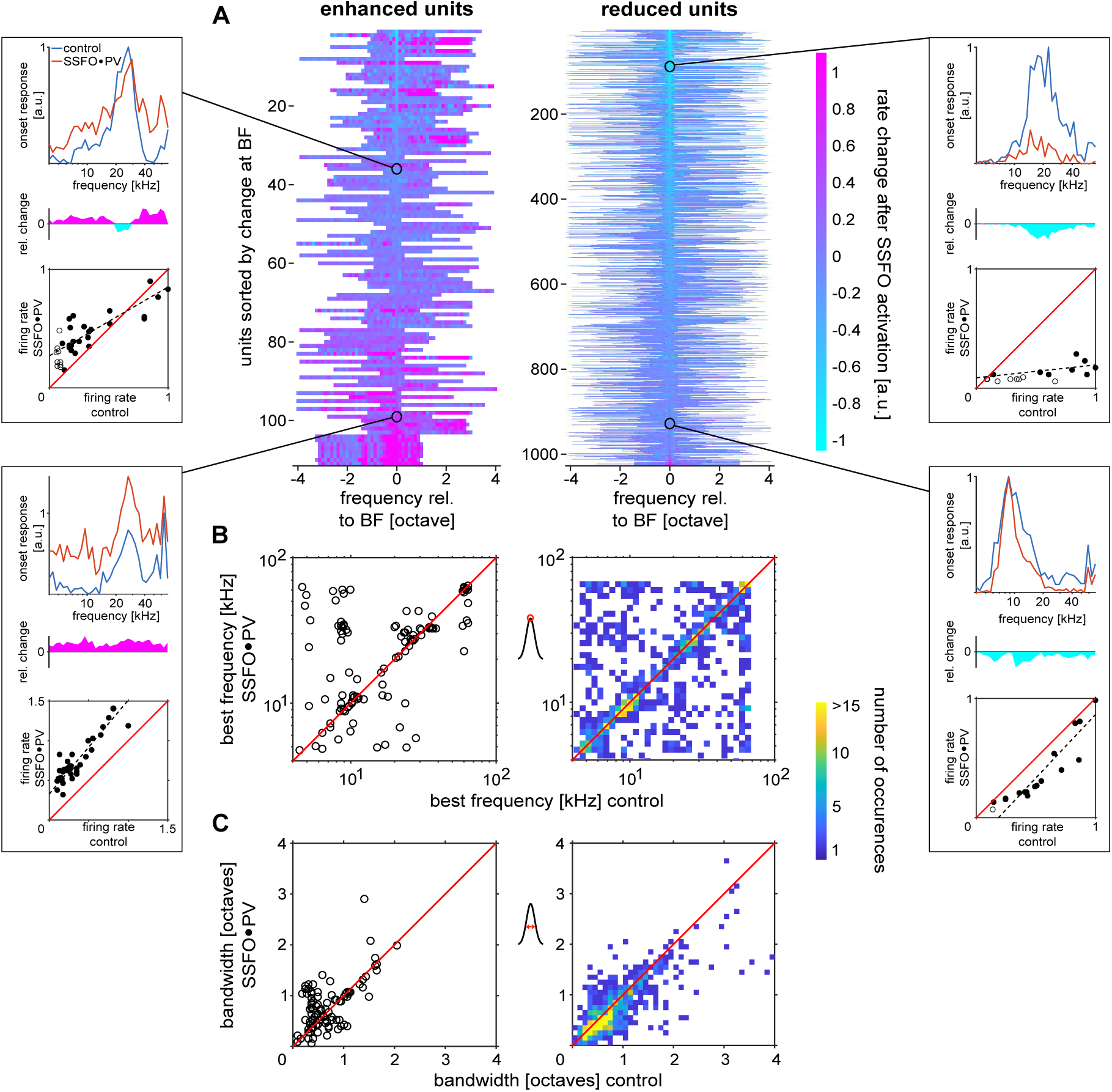
Sustained activation of PV+ cells has minimal effects on tuning best frequency and bandwidth. Tuning curves were generated for each unit from the mean firing in a short window after stimulus onset (15-55 ms relative to sound onset). (**A**): Overview of tuning curve changes during SSFO•PV trials, grouped by the mean effect the manipulation had on the unit (left: enhanced units, n = 111 units; right: reduced, n = 1027 units). Within each group, units are sorted by the amount of change in SSFO•PV trials at their best frequency (BF): units with a large decrease in the response (light blue) at BF are located at the top, units with a large increase (purple) at BF are located at the bottom (examples boxes). Boxes at the side: tuning curves of example units for control condition (blue) and during SSFO•PV trials (red); relative change in the tuning curves (center); and difference in firing rate between the two conditions (bottom). The black dashed line marks the result of a major axis regression between control and SSFO•PV trials. To avoid threshold effects, only frequencies with a firing rate greater than 10% of the maximum rate in both conditions were used for the major axis regression (filled circles). Black circle and line indicates which unit in A is shown in the box. (**B**): Best frequency during SSFO•PV vs. control trials (left: enhanced units, n = 111 units; right: reduced units, n = 1027 units). (**C**): Bandwidth (measured at 50% of peak amplitude) for SSFO•PV trials plotted against the bandwidth for control trials (left, n = 93 units; right, n = 876 units). Because of the large number of enhanced units (right panels in B and C), data from these was plotted as histograms, while data for reduced units is presented in scatter plots.

A selective reduction at off-BF frequencies would result in narrower overall tuning as measured by the bandwidth (BW). Consequently, we found a small but significant reduction of BW in the reduced group (Fig. 3C, right panel, median change in bandwidth 0.06 octaves, p<10^−25^, Wilcoxon sign-rank test). For enhanced units, we found a significant (p=0.0396, Wilcoxon sign-rank test) slightly small median increase in bandwidth (10^−15^ octave).

While BW was slightly changed, activation of PV+ cells did not affect the units’ best frequency tuning (Fig. 3B, median change in frequency in each group = 0 octave).

Thus, despite a considerable reduction of the overall firing rate during sustained PV+ activation (Fig. 1), both BF and BW where well conserved during SSFO•PV trials, with BW changes of less then a semitone on average and almost no change in BF (Fig. 3).

### Reduction of firing is predominantly divisive

Since we observed very little change in neuronal frequency tuning, we hypothesised that sustained activation of PV+ interneurons may provide a means for the context-dependent modulation of divisive gain control. Divisive action on tuning curves should result in little or no change in tuning characteristics due to the firing-rate-dependent adjustment for each frequency in the tuning curve. Conversely, previous studies using short-term optogenetic stimulation have reported that PV+ activation results in a mixture of divisive and subtractive changes (Seybold et al. 2015; Phillips and Hasenstaub 2016). Here we asked whether sustained activation would also result in mixed effects or provide means for predominantly divisive modulation.

We calculated the tuning curves for each unit for both control and SSFO•PV trials and normalised them to the peak of the control tuning curve. We then compared the normalised firing rate per frequency for tuning curves from SSFO•PV trials to those from control trials using linear regression (see Fig. 3, example boxes, bottom). For further analysis, we excluded tuning curves with a correlation coefficient smaller than 0.5 (Fig. 4, A, left). In order to describe patterns of modulation in the data, we extracted the slope and the y-intercept from the fit. If there were no difference between the tuning curves in the control condition and after PV+ activation, the slope would be 1 while the y-intercept would be 0 (Fig. 4, A, red vertical lines).

**Figure 4.**
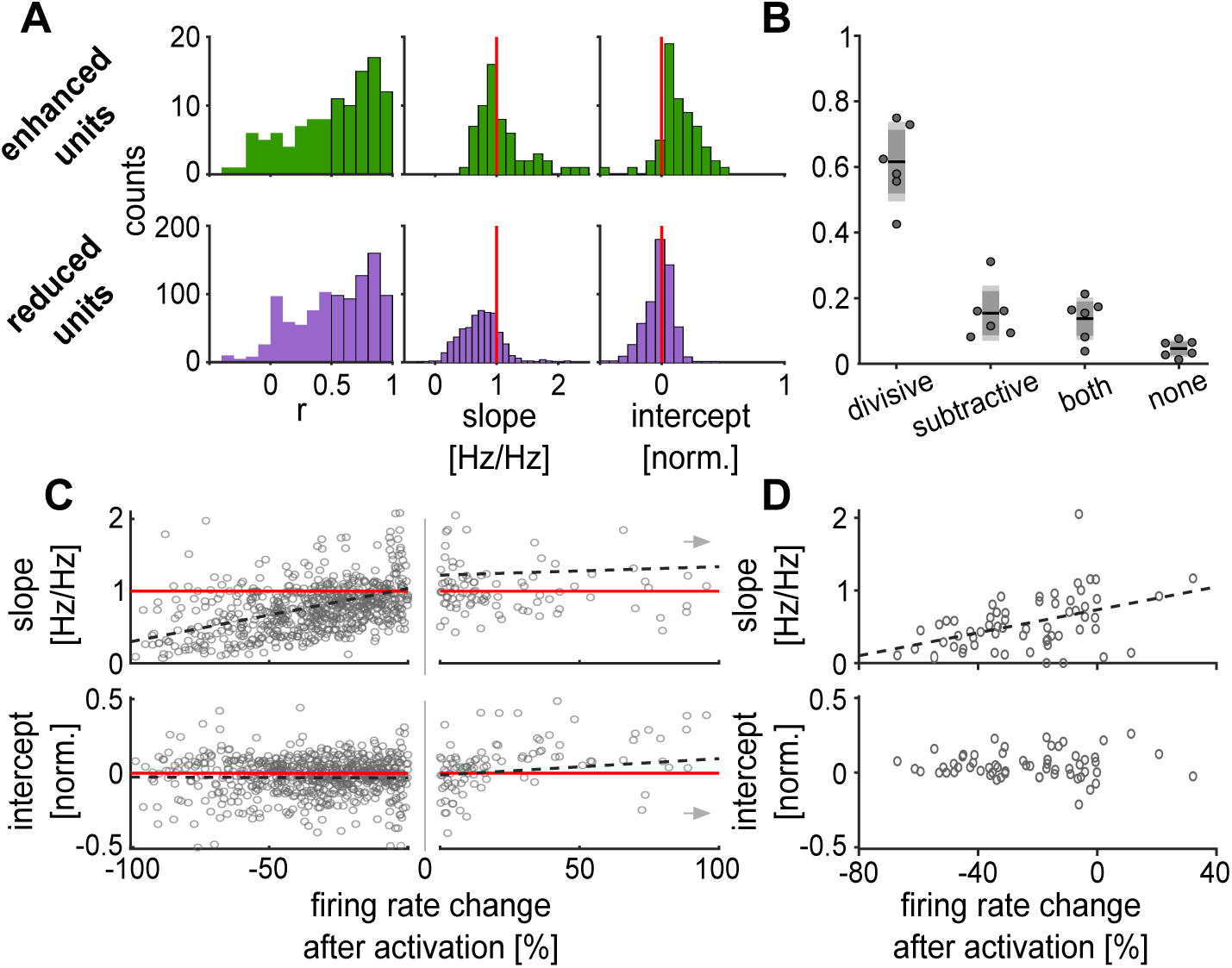
Sustained activation of PV+ cells results in a predominantly divisive change in firing rates. (**A**): Distribution of parameters of the linear fits between tuning curve in control and SSFO•PV trials (Fig. 3 example boxes) for enhanced units (top, n =111) and reduced units (bottom, n = 1027). Units with a correlation coefficient r < 0.5 (left) were excluded from further analysis of the fit. The red line indicates no change in the slope (center) or shift of the y-intercept (right). For illustrative purposes, the intercept of one enhanced unit is not displayed (intercept = − 2.36). (**B**): Proportion of units with significant divisive and/or subtractive effects in SSFO•PV trials compared to control (mean ± SEM (light grey) and ± standard deviation (dark grey). Circles represent the proportion in each individual animal (n = 7). (**C**): Relationship between firing rate change in each individual unit and the slope (top) and intercept (bottom) parameters of the linear fits. Dashed lines represent linear regressions, separately fit for reduced (n = 694) and enhanced (n = 110) units. For illustrative purposes, units with a change > 100% are not displayed (grey arrows), but were included in the fit. (**D**): Same as in (C) for the median rate change and median fitted parameters of all units recorded in each recording position (n = 61).

The reduced units displayed very little subtractive change based on the y-intercept (Fig. 4A, right, median (IQR) = −0.01 (0.141)), but a clearly reduced slope (Fig. 4A, center, median (IQR) = 0.75 (0.41)), indicative of mostly divisive gain control (Fig. 4A, bottom, n = 581). However, for enhanced units, the tuning curves were mostly additively shifted upwards (right, median (IQR) = 0.11 (0.2)), and revealed a broader distribution for slope changes with a small multiplicative effect on average only (center, median (IQR) = 0.99 (0.53)).

In order to quantify whether the effect of sustained PV+ activation on individual units was divisive or subtractive, we categorized the units according to whether the regression slope and y-intercept differed significantly from 1 and 0 respectively (Fig. 4B, n = 7). In all tested animals, significant divisive suppression was clearly the dominant effect (proportion of purely divisive units: mean (SD) = 0.6166 (± 0.1212), SEM = 0.0495). In contrast, we observed only few units with a subtractive suppression (mean (SD) = 0.15434 (± 0.0838), SEM = 0.0342), a combination of both (mean (SD) = 0.1378 (± 0.0648), SEM = 0.0264), or neither (mean (SD) = 0.0464 (± 0.0253), SEM = 0.0103).

For individual units, overall reduction in firing rate could be well explained by a reduction of slope (Fig. 4C, top, correlation coefficient r = 0.34, p = 1.2e-19, n = 694). On the other hand, we did not find a relationship between rate changes and y-intercepts for individual units (bottom, correlation coefficient r = −0.01, p = 0.847, n = 694).

It has been suggested that the relative strength of divisive and subtractive action mediated by PV+ cell activation could be related to the extent of activation (Seybold et al. 2012; Lee et al. 2014). As an indirect measure of PV+ cell activation, we compared median firing rate change within single experiments to the median slope and y-intercept by pooling all recorded units at the respective position (Fig. 4D, n = 61). The results of this analysis confirmed our finding from the individual units; we again observed a high correlation of the slope and mean rate reduction (top, correlation coefficient r = 0.45, p = 3e-4) and a mostly constant intercept (bottom, correlation coefficient r = 0.01, p = 0.93). Thus, most suppression of firing is mediated by divisive changes. Moreover, division scaled linearly with the amount of suppression, indicating that sustained activation of PV+ units results in a divisive scaling of neuronal output, and depends on the strength of inhibitory drive.

### Spectro-temporal receptive fields are divisively scaled and their structure conserved

The prominent divisive effect of sustained PV+ activation on tuning curves with only a minor impact on tuning best frequency and bandwidth led us to the question of how such effects would generalise to more complex stimuli. Therefore, we investigated the impact of SSFO manipulation of PV+ activity on spectro-temporal receptive fields (STRFs) estimated from responses of cortical cells to dynamic random chord (DRC) stimuli.

We found that many STRFs estimated from DRC responses were divisively or multiplicatively modulated for reduced and enhanced cells respectively, which in turn meant that DRC responses predicted from the STRFs were divisively suppressed or multiplicatively enhanced (Fig. 5A).

**Figure 5.**
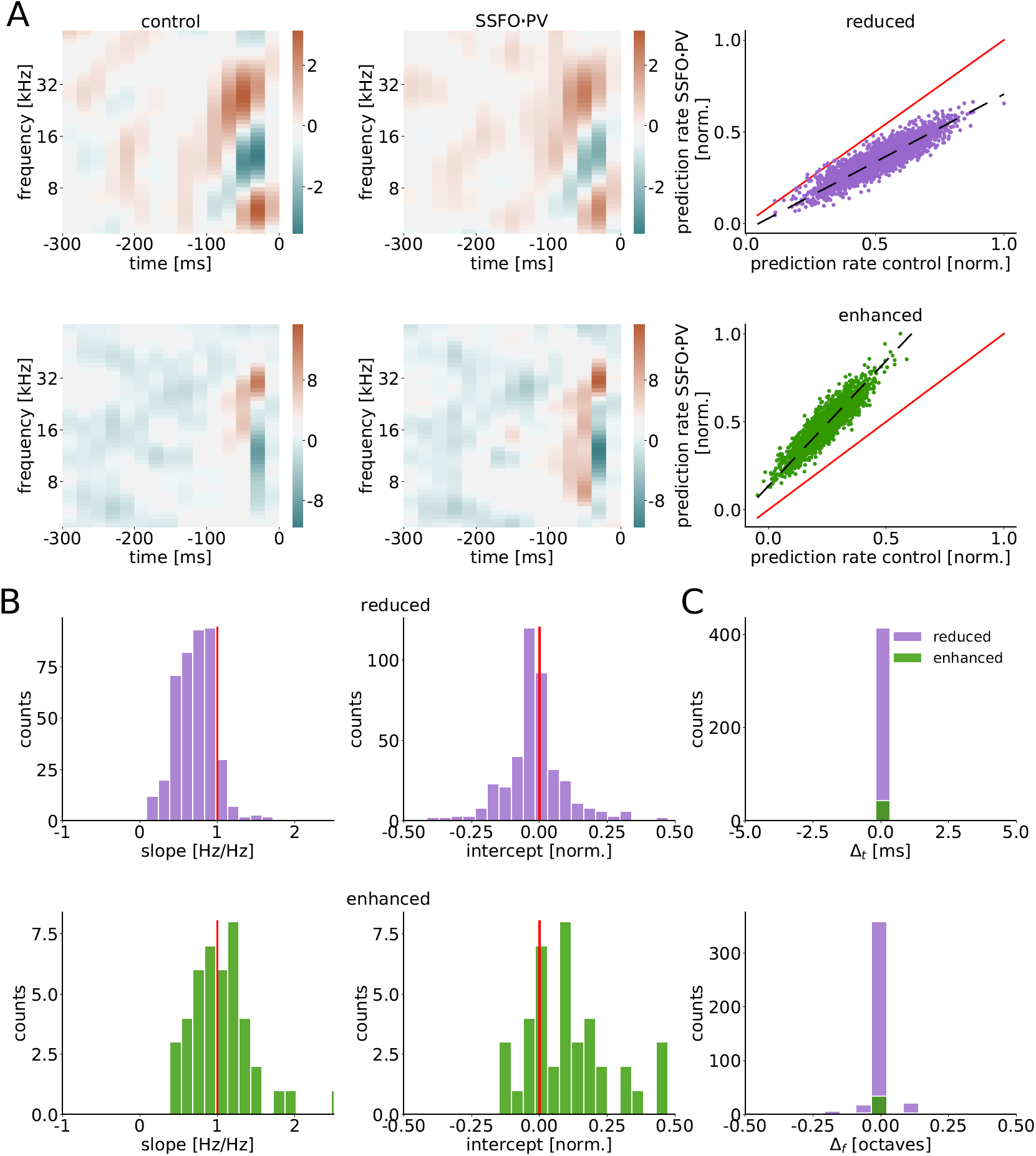
Spectro-temporal receptive fields are divisively scaled by sustained PV+ activation, but their structure is preserved. (**A**): STRF for a reduced unit (upper) and an enhanced unit (lower) in control and PV-activated conditions. The STRF predicted activity for the reduced unit is divisively modulated in PV-activated compared to control conditions. The STRF-predicted activity for the enhanced unit shows a multiplicative enhancement. (**B**): For reduced (top) and enhanced (bottom) units, slopes (left) and intercepts (right) of linear fits to STRF predicted activities in PV-activated versus control conditions. Only units with a correlation coefficient r > 0.5 are included (enhanced units n = 44 and reduced units n = 419). We note that a small fraction of units (1 to 2% of reduced units, and 5 to 9% of enhanced units) lie outside the plotted bounds and are excluded from the histograms to enable better visualisation of the bulk of the data. (**C**): For most units, sustained PV-activation does not shift the STRF in time and frequency. 1% of reduced units and 2 to 5% of enhanced units lie outside the plotted bounds and are excluded for better visualisation of the rest.

Linear fits to the STRF predictions for PV-activated versus control conditions show that most reduced cells were strongly modulated through the slope, i.e. divisively (Fig. 5B top; slope median (IQR) = 0.72 (0.33) and y-intercept median (IQR) = −0.02 (0.09)). In contrast, enhanced cells showed more heterogeneous effects, with some units exhibiting strong modulation through the intercept, i.e. additively (Fig. 5B bottom; slope median (IQR) = 1.01 (0.44) and y-intercept median (IQR) = 0.09 (0.20)). However, as previously explained, for the majority of cells both DRC responses and single-tone responses were reduced, not enhanced, by sustained PV+ activation (Fig. 1D). Overall, therefore, the STRF analysis of DRC responses reveals predominantly divisive effects of sustained PV+ activation, in agreement with analysis of single-tone responses. We note that restricting the analysis to the subset of units with the highest coefficients of determination of the estimated STRFs confirms these results (Supplementary Fig. S1).

We also found that for the majority of cells, sustained PV+ activation did not shift the STRF in time and frequency (Fig. 5C). We quantified this by computing the lag in time and frequency that maximised the cross-correlation between the STRFs in the PV-activated and control conditions. These time and frequency lags were tightly concentrated near 0 ms and 0 octaves (Fig. 5C; for both reduced and enhanced units, time lag median (IQR) = 0.0 (0.0) ms and frequency lag median (IQR) = 0.0 (0.0) octaves). Thus, the minor impact of sustained PV+ activation on the best frequency and bandwidth of responses to pure tones generalised to a minimal impact of the manipulation on spectrotemporal tuning.

### Divisive scaling generalizes to naturalistic stimuli

We found consistently divisive scaling of single-unit responses both for ST (Fig. 3) and DRC stimuli (Fig. 5), conserving the receptive fields of the units. We next asked whether divisive scaling would transfer to more naturalistic stimuli. To address this question, we recorded responses in a subset of the animals (n = 4) to a set of animal vocalizations varying in temporal and spectral structure, and applied the same manipulation as for ST and DRC stimuli. We recorded from a total of 513 responsive units in all three paradigms.

Many units locked their spiking activity to the envelope of the natural stimuli (Fig. 6A). During SSFO•PV trials, responses were similar and typically a scaled version of the responses in the control trials (Figs. 6B,C, left panels). When we compared spike rate modulation in all three paradigms, we observed highly conserved divisive scaling in the majority of units. Units that were enhanced in one paradigm, also were enhanced in the other two (Fig. 6B), and units with decreased activity during SSFO•PV trials scaled divisively in all three paradigms (Fig. 6C).

**Figure 6.**
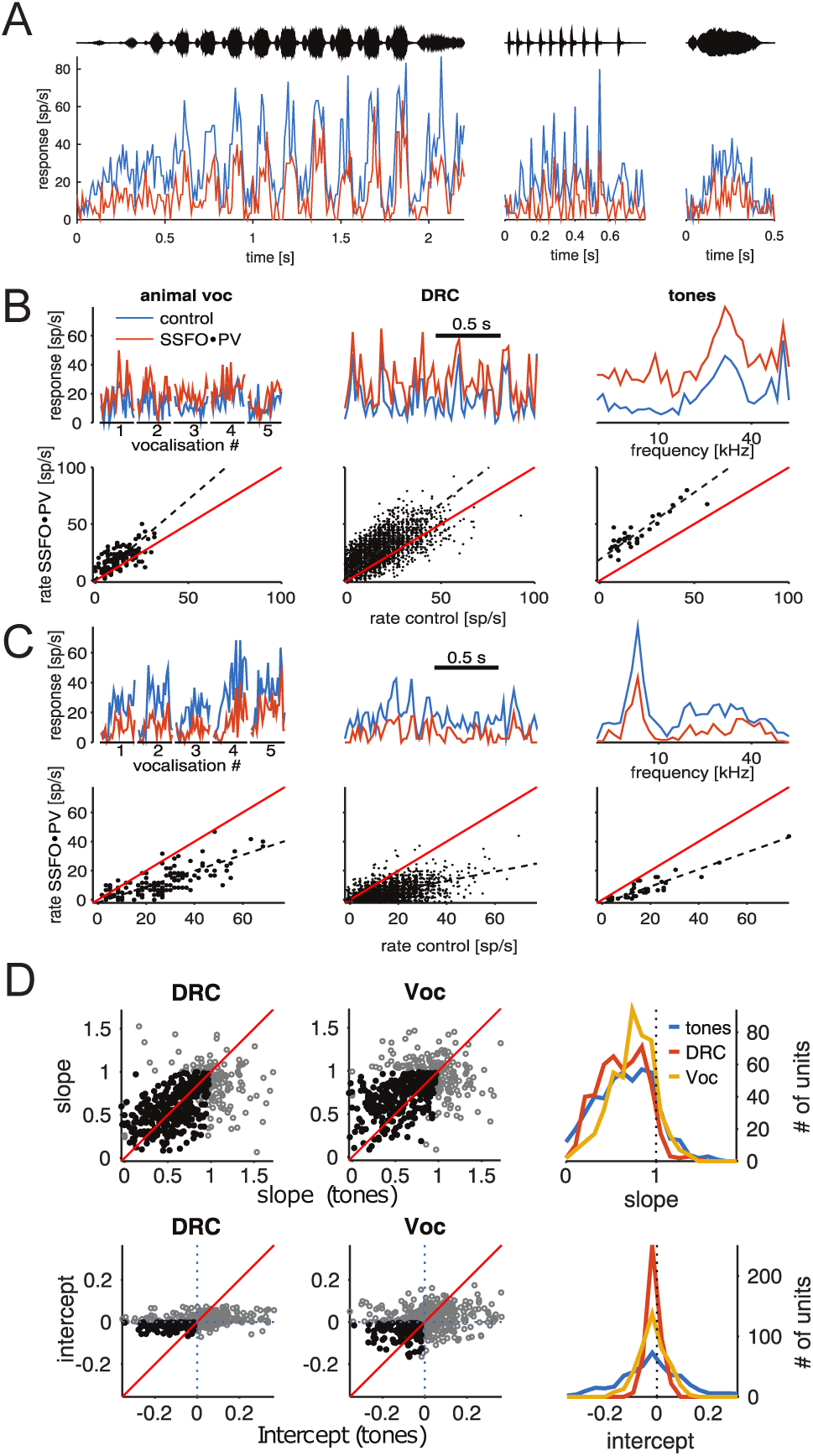
Divisive action generalizes to naturalistic stimuli and is consistent across stimulus paradigms. (**A**): Example responses to three different animal vocalizations. On the top, the time course of the respective vocalization is depicted (vocalizations 1-3 in panels B and C). Blue line: control, red line: SSFO•PV. (**B**): Effect of sustained PV-activation on the responses of one enhanced unit to all three different paradigms (from a population of 513 cells with recordings in all three paradigms). Top left: Animal vocalizations – 500ms snippets from all five vocalizations used. Top center: DRC – 2 second snippets taken from continuous DRC stimulation. Top right: Tones – tuning curves obtained from tone onsets (see Fig. 3). Lower panels show comparison of firing rate in control and SSFO•PV trials in the respective paradigm, including the full recording. Dashed line depicts the linear fit. (**C**): Same as B, but for one unit with decreased firing rates in the SSFO•PV condition. (**D**): Relationship between linear fit parameters observed in different stimulus paradigms. Scatter plots depict slope and intercept fits for single units, comparing tones and DRC stimulation (left) or tones and vocalization stimuli (center). Grey, open markers are units for which the respective linear fit parameter was not significantly different from what would be expected if PV-activation had no effect (slope=1, intercept=0); filled, black markers are those with parameter value significantly different from the null condition (one sided, *p* < 0.05). Right, histograms of fit parameter values for all three stimulus paradigms.

This was confirmed when we looked at divisive and subtractive changes in SSFO•PV trials compared to control (Fig. 6D). Neither slopes nor intercepts changed between the ST and DRC paradigms in single units (Fig. 6D). Slopes were slightly less reduced for the vocalizations, but still highly correlated for single units (Fig. 6D, upper panels). The medians of the intercepts were close to zero for all paradigms, with a much larger variance in estimates from the ST paradigm than from DRCs and vocalizations (Fig. 6D, lower panels).

### Recurrent network model with power-law input-output functions captures divisive modulation and conservation of tuning

As shown above, in most auditory cortical neurons and across stimulation paradigms, sustained PV+ activation produces divisive modulation of auditory responses, with conservation of response properties such as tuning best frequency and STRF structure. These observations suggest that a unifying mechanism might be at play.

Previous work has shown that responses of single neurons in V1 are captured by power-law input-output functions (Priebe and Ferster 2008), and that models of recurrent excitatory-inhibitory networks with such input-output functions can show divisiveness in response to the activation of the inhibitory population (Litwin-Kumar et al. 2016). Here we illustrate that this same mechanism captures the main findings reported here.

We assume that the recorded cortical cells are part of an excitatory-inhibitory network, where neurons have power-law input-output functions, and inherit their bell-shaped tuning from the thalamic input (Fig. 7A). Optogenetic activation of inhibitory (PV+) neurons is modelled as an additive current, heterogeneous across inhibitory cells. Such heterogeneity is introduced to account for the likely varied genetic expression of SSFO and diverse light exposure across PV+ cells. As expected, the network model leads to divisive modulation of excitatory cell tuning (Fig. 7B-D). Furthermore, inhibitory cells show a mixture of effects, from divisive to additive and multiplicative, depending on the magnitude of the direct optogenetic current; responses of inhibitory cells with low direct activation are typically reduced overall, as their net recurrent input is decreased, while responses of inhibitory cells with higher direct activation are enhanced, as their net input is increased. Overall, across the whole network, optogenetic modulation leads mostly to divisive modulation of tuning curves, with a small fraction of cells being subtractively or both subtractively and divisively modulated (Fig. 7D-E). We note that heterogeneity in the optogenetic activation leads to some inhibitory cells being reduced in addition to the excitatory cells, which could explain the absence of a clear separation between the spike shapes of the enhanced and reduced units in our electrophysiological recordings (Fig. 2). Finally, our model shows that for the large majority of neurons, there are minimal or no changes in tuning best frequency and bandwidth between conditions, in agreement with our experimental data (Fig. 7F and Fig. 3B,C).

**Figure 7.**
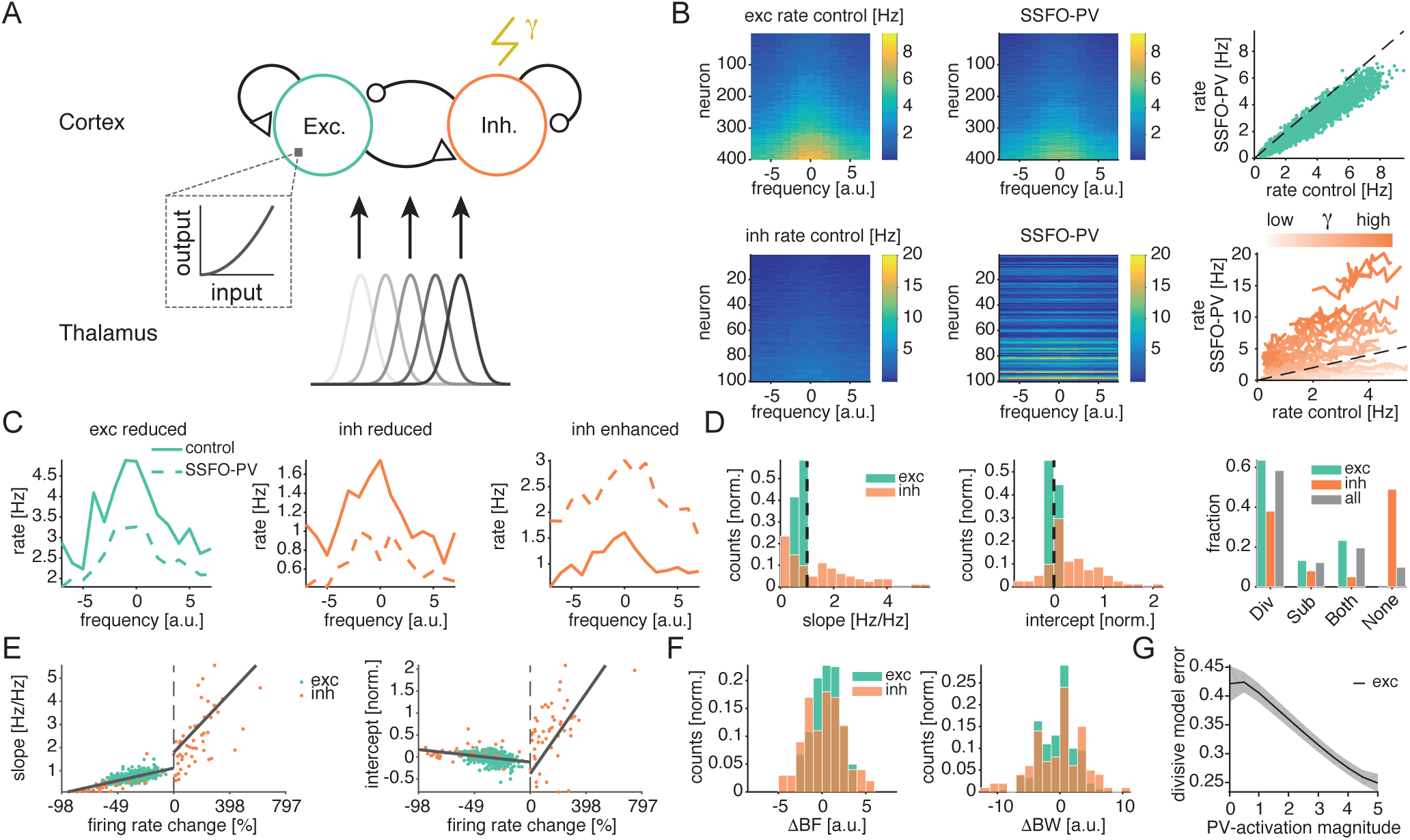
Network model with power-law input-output functions captures divisive modulation of responses and preservation of tuning. (**A**): Schematic of network model. Recorded cortical cells are modelled as an excitatory-inhibitory network with power-law input-output neural functions. Optogenetic activation of inhibitory (PV+) neurons is modelled as an additive current, heterogeneous across inhibitory cells. Bell-shaped frequency tuning in cortical units is inherited from the thalamic input. (**B**): Model tuning curves of excitatory (upper) and inhibitory (lower) cells in control and PV-activated conditions. Excitatory neural activity is typically divisively modulated in PV-activated compared to control conditions. Inhibitory cell activity shows a broad range of effects, i.e. divisive, additive and multiplicative effects, depending on the magnitude of direct PV-activation. (**C**): Three representative tuning curves in control and PV-activated conditions: excitatory (left) and inhibitory (middle) cells divisively reduced during PV-activation, and inhibitory cell (right) enhanced during PV-activation. (**D**): (Left and middle) Slopes and intercepts of linear fits to tuning activities in PV-activated versus control conditions. We note that 15% to 20% of the units lie outside the plotted bounds and are excluded to enable better visualisation of the bulk of the data. (Right) Most cells are divisively modulated during sustained PV-activation. (**E**): (Left) Slope is positively correlated with change in firing rate (r = 0.72 and p = 1.40e-71, in the domain of negative rate changes; r = 0.52 and p = 2.43e-05, in the domain of positive rate changes). In contrast, (right) intercept is negatively correlated with negative rate changes (r = −0.35, p = 2.19e-14) and positively correlated with positive rate changes (r = 0.50, p = 5.11e-05). (**F**): Differences between PV-activated and control conditions of tuning best frequencies (left, ΔBF) and tuning bandwidths (right, ΔBW) are typically close to zero. (**G**): Prediction. Mean error, and respective SEM, of purely divisive model (intercept at zero) for the activity of excitatory units modulated by sustained PV+ activation, as a function of PV-activation magnitude (including zero), for multiple noise instances.

In summary, a recurrent excitatory-inhibitory network with power-law input-output functions captures our main experimental findings— particularly the observation that the dominant effect of sustained PV+ activation is divisive modulation of firing rate with conservation of response properties such as tuning best frequency and bandwidth. Moreover, as illustrated in Supplementary Fig. S2, both recurrent connectivity and power-law input-output functions contribute to the predominance of divisive effects of PV+ activation. In the model, the impact of sustained PV+ activation on feedforward inputs is subtractive by design. However, synaptic inputs to model units (arising from both feedforward and recurrent connections) can be either subtractively or divisively modulated, and firing rate outputs of model units (arising from feedforward and recurrent inputs and power-law input-output functions) are often more strongly divisive modulated than their inputs (Supplementary Fig. S2). Thus both recurrent connectivity and power-law input-output functions contribute to strengthening divisive modulation by sustained PV+ activation.

Lastly, the model predicts that the stronger the activation of PV+ interneurons, the better a purely divisive model (intercept constrained at zero) explains the modulation of excitatory units with PV+ activation (Fig. 7G). This finding indicates that the lower proportion of divisive effects observed in Seybold et al. 2015 could be due to a weaker PV+ activation compared to the one applied in our study (Fig. 7G). More generally, the prediction suggests that PV+ interneurons could be used as a flexible modulator of divisiveness in the network.

## Discussion

Here we asked whether sustained activation of PV+ cells on slower time scales than tested before could serve as a general mechanism for implementing divisive gain control. To this end, we used a bi-stable variant of Channelrhodopsin (SSFO) to activate and deactivate PV+ cells on the multi-second time scales that are typical for contextual modulation of PV+ cells (Kawaguchi and Shindou 1998; Metherate et al. 1992; Alitto and Dan 2012; Schneider et al. 2014; Petrie et al. 2005). This also enabled us to test effects of PV+ activation not only on brief sounds such as single tones (ST), but also on prolonged sounds including complex natural stimuli (NS) and dynamic random chords (DRC). Our results show that sustained activation of PV+ interneurons produces divisive modulation of cortical responses (Fig. 3) which is consistent across trials (Fig. 1), single units, electrode positions (Fig. 4), and greatly differing stimulus paradigms (Fig. 6). Furthermore the divisive change preserves the main response properties of cortical units in both the spectral (tuning curves Fig. 3; receptive fields Fig. 5) and temporal domains (Fig. 5). Finally, we were able to capture the experimental findings in a recurrent network model with power-law input-output neuronal functions (Fig. 7).

### Cell-to-cell variability in effects of sustained PV+ activation

Although sustained PV+ activation produced strong suppression of neural responses in a large majority of neurons (“reduced units”), a subset of recorded cells instead showed enhancement of firing rate during SSFO•PV trials (“enhanced units”) (Fig. 1). The distribution of modulatory effects was essentially unimodal across the recorded population, with no clear separation between reduced and enhanced units. However, in individual units, modulatory effects were reproducible and consistent across different auditory stimuli, and at the population level, a similar diversity of modulatory effects was observed in different experimental animals. Moreover, diverse effects of sustained PV+ activation were observed even among neighbouring neurons at a single electrode position within individual mice. These observations suggest that cell-to-cell variability in effects of sustained PV+ activation might arise not only from differences in viral expression or spread of light at different electrode positions and in different mice, but also from locally heterogeneous synaptic strength of (PV+) inhibitory connections or locally heterogeneous activation of PV+ cells. Units with strong PV+ inputs could be more strongly affected by the activation of PV+ cells than neighbouring units with weak PV+ inputs. Such local heterogeneity in effects of sustained PV+ activation was implemented in our model, and reproduced the overall variability of effects observed in the data. However, inter-animal, inter-position and intra-position sources of variation could not be differentiated from our data.

For cells with reduced activity during sustained PV+ activation, the extent of reduction was associated solely with the strength of division (Fig. 4). In contrast, cells with enhanced firing rates predominantly showed an additive change in their responses (Fig. 4A). It is conceivable that these units were primarily PV+ cells that had been directly activated. This interpretation is consistent with the observation that spike waveforms for enhanced units had narrower peaks and deeper troughs than for reduced units (Fig. 2). If PV+ units were enhanced and non-PV+ units reduced in firing rate, why did we not observe a clear bimodal distribution of firing rate changes? It has been shown that activation of PV+ cells in some layers can functionally disinhibit PV+ cells in other layers (Moore et al. 2018). These indirect disinhibitory effects, along with variable activation of PV+ cells and variability in PV+ to non-PV+ cell connections, could produce high variation in effective firing rate changes in PV+ units. This hypothesis is supported by results from the network model, where we implemented heterogeneous activation of PV+ cells and also observed a continuous distribution of effects of sustained PV+ activation on the tuning curves for inhibitory neurons.

### Comparison to previous findings on optogenetic PV+ manipulation in AC

In comparison to previous studies on the computational roles of cortical inhibitory interneurons in the auditory cortex (e.g., Seybold et al. 2015; Phillips and Hasenstaub 2016), we found a more divisive and less broadly mixed effect of PV+ activation. Our optogenetic approach differed most significantly from previous work in the timing of the optogenetic manipulation.

While other studies have combined standard ChR2 and stimulus-synchronized light activation, the SSFO variant used here allows for decoupling of light activation and sensory stimuli. It has already been shown for visual cortex that the effect of PV+ activation depends on the relative timing of the circuit manipulation and the sensory stimulus (Atallah et al. 2012; Lee et al. 2012; Wilson et al. 2012; Lee et al. 2014). Similar factors may explain some of the differences between our results and previous findings (Seybold et al. 2015; Phillips and Hasenstaub 2016). Both transient, stimulus-locked activation and slowly varying, sustained activation may be important for contextual processing and top-down control. While modulatory effects on PV+ cell activity related to locomotion (Schneider et al. 2014; Polack et al. 2013; Pakan et al. 2016) and vigilance (Kawaguchi and Shindou 1998; Kuchibhotla et al. 2016) typically last for several seconds to minutes, some forms of neuromodulation may also include fast, transient components at the time scale of tens of milliseconds (Muñoz and Rudy 2014; Poorthuis et al. 2014; McGinley et al. 2015). An exact description of time scales for contextual modulation of sensory computation may be crucial for a detailed understanding of these processes (Edeline 2012).

A second technical aspect that differed between our study and previous related work was the location of the optogenetic stimulation site within the auditory cortex. A study using local stimulation of PV+ cells in different layers showed that activation in one layer may have deactivating effects on PV+ cells in other layers and vice-versa (Moore et al. 2018). Other studies stimulated at the surface of the cortex using low light levels (< 0.5mW/mm^2^ (Seybold et al. 2015; Phillips and Hasenstaub 2016; Aizenberg et al. 2015). Since light transmission drops in brain tissue by 50% every 200 µm (Yizhar et al. 2011), surface illumination may have activated mainly PV+ cells in superficial cortical layers, potentially mostly Chandelier cells with large arborizations in layer 1 (Inan and Anderson 2014). In contrast, our fibre tip was typically placed in middle layers (IV and adjacent) and oriented tangentially to the cortical surface, probably mostly activating PV+ cells throughout middle layers. These layers contain more PV+ cells than layer I and predominantly Basket rather than Chandelier cells (Miyamae et al. 2017).. Chandelier and Basket cells synapse onto different segmental areas of their target cells, and may thus differ in their impact on synaptic integration in the post-synaptic cell (Markram et al. 2004). Activating different proportions of these distinct PV+ cell types could well result in different proportions of subtractive (Chandelier cells, axonal targeting) and divisive (Basket cells, pre-axonal targeting) firing rate changes, further explaining some differences between our results and previous findings. Additionally, our network modelling suggests another explanation for the higher fraction of divisive effects on our dataset: independently of the targeted PV+ cell type, higher light levels reaching PV+ cells and the resulting higher magnitude of PV+ activation is sufficient to explain a larger proportion of divisive effects, a direct consequence of the power-law nature of the input-output functions (Fig. 7G).

### Use of bi-stable optogenetic tools for probing cortical inhibition

Standard optogenetic tools allow for manipulation of cell activity during laser illumination only, and the duration of illumination is limited to a few hundred milliseconds by the potential for photodamage and temperature increase (Yizhar et al. 2011). This limitation means that most optogenetic manipulations of sensory processing are performed using short and low-complexity stimuli. The SSFO variant we used here allowed us to manipulate PV+ cells during complex and naturalistic stimuli and to explore how effects of PV+ activation generalize across stimulus sets and over time.

In addition, bi-stable optogenetic tools as we used here may be an important means of understanding the effects of neuromodulation of cortical circuitry (Edeline 2012) mediated by differential activation of distinct groups of interneurons at slower time scales(Paul et al. 2017). Several neuromodulators have been shown to specifically affect cortical PV+ cells (e.g. serotonin (Puig et al. 2010), acetylcholine (Metherate et al. 1992; Alitto and Dan 2012), norepinephrine (Toussay et al. 2013)) on time scales of seconds to minutes. Thus, using SSFO enabled us not only to implement complex and prolonged auditory stimuli and to avoid laser onset and offset artefacts, but also to attain an activation of PV+ cells at new and physiologically relevant time scales.

### Relation to previous modelling work on inhibition in auditory cortex

As discussed above, recent optogenetic studies in the auditory cortex have provided new clues about the role of inhibition in shaping the dynamics of cortical networks. Several theoretical models have been put forward to account for diverse datasets, from models with synaptic depression (Loebel et al. 2007; Aizenberg et al. 2015), to inhibition-stabilised networks (Moore et al. 2018) and feedforward models (Seybold et al. 2015). Here, we provided a minimal model – a recurrent network model with power-law input-output functions and heterogeneous optogenetic activation – that can account for three main features of the experimental data: the prominence of divisive effects of PV+ activation, the minimal impact of PV+ activation on the tuning BF and BW, and the absence of a clear separation of spike shapes between reduced and enhanced units. Furthermore, the model reconciles our results with the previous conflicting conclusion that PV activation leads to a lower fraction of divisive effects (Seybold et al. 2015): in our model, this decreased fraction can be explained by a decrease of the magnitude of PV activation (Fig. 7G).

The effects reported by Seybold et al. 2015 and Phillips and Hasenstaub 2016 were well-replicated by an abstract feedforward model (Seybold et al. 2015), perhaps in part because effects of transient optogenetic activation time-locked to short tonal stimuli are dominated by feedforward connectivity. In contrast, network modulation in general (and sustained PV+ activation in particular) may engage network dynamics more fully and thus might be better explained by models with recurrent dynamics.

In our model, the divisive modulation by PV+ activation is a consequence of both power-law neuronal input-output functions and recurrent dynamics. This is demonstrated by the following observations: first, in some network units, the synaptic inputs (arising from feedforward and recurrent connections) are subtractively modulated by sustained PV+ activation while output firing rates (shaped by feedforward and recurrent inputs and power-law input-output functions) are strongly divisively modulated; in contrast, in other model units, both the synaptic inputs and the firing rate outputs are divisively modulated, even though the feedforward inputs alone would be subtractively modulated (Supplementary Fig. S2). Therefore both recurrent connectivity and power-law input-output functions contribute to the predominance of divisive modulation in the model outputs.

Given its recurrent nature, our model could be a building block for improving understanding of the full complexity of neuromodulatory effects, which are known to engage recurrent network dynamics (Edeline 2012) as well as other interneuron classes (e.g. SOM+ and VIP+ cells).

### Functional implications for cortical computation

Divisive gain control has been proposed to be one of the canonical neural computations performed by cortical circuits (Salinas and Thier 2000; Carandini and Heeger 2011). A range of computationally important functions in sensory processing are attributed to divisive gain control, including adaptation to stimulus statistics (Rabinowitz et al. 2011), invariant stimulus encoding (Rabinowitz et al. 2013), optimised sensory discrimination (Guo et al. 2017), foreground-background separation (Busse et al. 2009), and selective attention (Ruff and Cohen 2017). Most of these functions depend on behavioral and sensory context and therefore need to be under control of modulatory mechanisms. We show here that prolonged, low-level activation of PV+ interneurons provides a means for flexibly modulating such gain control in auditory cortex. PV+ neurons are the target of neuromodulators such as serotonin (Puig et al. 2010) and acetylcholine (Metherate et al. 1992; Alitto and Dan 2012), and are differentially activated in specific behavioral states such as task engagement (Kuchibhotla et al. 2016) and locomotion (Schneider et al. 2014; Zhou et al. 2014), putting them in an optimal position to mediate necessary changes in sensory processing at the time scale of seconds to minutes. Our results demonstrate that sustained activation of PV+ interneurons in auditory cortex on this time scale produces robust divisive gain control, not only for brief tones but also for continuous, complex and natural stimuli.

## Methods

### Subjects

All data presented here were obtained from eight male B6.Cast/PVALB-Cre mice. This line is a cross between B6.CAST-Cdh23 (Stock number 002756, Jackson Laboratory, Bar Harbor, ME, US) and PVALB-IRES-Cre mice (Stock number 008069, Jackson Laboratory, Bar Harbor, ME, US), crossed and reared in the animal facilities of the University of Oldenburg. These animals are not susceptible to developing the early-onset age-related hearing loss typical of the standard C57BL/6 strain (Johnson et al. 1997). Cre-recombinase was expressed in inhibitory parvalbumin-positive (PV+) neurons (Hippenmeyer et al. 2005), enabling us to manipulate those neurons optogenetically following injection of the Cre-dependent viral vector. All mice were housed separately following surgical implantation with recording devices and maintained on a reversed 12h/12h light-dark cycle at approx. 23 °C with access to free water and food.

All procedures were performed in accordance with the animal welfare regulations of Lower Saxony and approved by the local authorities (State Office for Consumer Protection and Food Safety / LAVES).

### Surgery

Mice were equipped with a chronic implant (probe) for optogenetics and recording at the age of 8 to 12 weeks. Animals were injected subcutaneously with 0.1 mg/kg meloxicam subcutaneously pre- and post-operatively to reduce pain and inflammation. During surgery, the animals were anaesthetized with 1.5 % isoflurane (initial: 2 %). Body temperature was monitored and held at approx. 37°C. Eyes were covered with ophthalmic ointment. Anaesthetized mice were fixed in a stereotaxic apparatus (Model 900, Kopf instruments, Tujunga, CA, US) with zygomatic bars. After skull exposure, a trepanation was drilled in the dorsal skull (relative to Bregma: anterior-posterior −2.6 mm, medial-lateral −2.9 mm, injection angle 24°, right hemisphere) to allow tangential access to the region designated AC in the Allen Mouse Brain Atlas (2011, www.mouse.brain-map.org). 1000 nl of adeno-associated virus (pAAV-EF1/a-DIOhChR2-(C128S/D156A)-EYFP, serotype 5, Vector Core, University of North Carolina, US) was injected (Nanofil 10 µl, WPI, Hertfordshire, UK) at 150 nl/min at each of two depths within auditory cortex (z=3.7 and z=3.9-4.0 mm from entry point in dorsal cortex). After virus injection, the probe was implanted with the electrode tips placed at the 3.7 mm depth initially. This placement resulted in a path of the electrodes that was tangential to the cortical surface at a depth of approximately 400µm, so most of our recordings likely stem from middle layers of core auditory cortex. In order to fix the implant microdrive onto the skull, five to six screws were drilled into the skull, one of which over the contra-lateral pre-frontal cortex served as reference for the recordings. The implant microdrive was then secured to the skull and screws with dental acrylic (Vertex SC, Vertex Dental, Zeist, Netherlands). After surgery, mice were given a recovery period between 7 and 14 days before recordings began.

### Implant design

Implants were custom-made at our laboratory. The implant consisted of eight twisted-wire tetrodes (17 µm, Platinum/10% Iridium California Fine Wire Company, Grover Beach, CA, US) concentrically arranged around a 105 µm optic fibre (FG105LCA Multimode Fiber, Thorlabs, Newton, NJ, US) for optogenetic manipulations and was attached to a microdrive (Axona, London, UK) to move the probe within AC. The electrode tips protruded 400 µm from the tip of the fibre.

### Optogenetic strategy

In the Cre-expressing PV+ cells of the mouse line, infection by the injected Cre-activated recombinant viral vector produced expression of a bi-stable ChR2 variant (Stable Step-Function Opsin, SSFO). Expression was confirmed via histological examination of brain tissue after the experiments. Expression spread through all layers of the auditory cortex and extended at least with a diameter of 1.5 mm radially in the medial-lateral and caudal-rostral dimensions. Efficiency of expression in PV+ cells was >95% (confirmed in three animals using imaging of immunostained PV+ cells and cells expressing the virally transmitted yellow fluorescent protein). SSFO was used to achieve prolonged, stable, low-level depolarization of PV+ cells (Yizhar et al. 2011), and has the advantage of its activation/deactivation interfering minimally with the timing of the sensory stimulus. Activation of SSFO was initialized by a pulse of blue light (447 nm, 2.5 mW for 2 s). SSFO cation channels remain open and induce a depolarizing input current until deactivation with a pulse of orange light (594 nm, duration 5 or 10 s at 2.5 mW).

### Electrophysiology

We recorded neural data extracellularly during auditory stimulation in alternating periods either without or with PV+ cell activation. These periods lasted each between 3 and 5 minutes. Continuous raw voltage traces were amplified and digitized using a 32-channel headstage (RHD2132, Intan Technologies, Los Angeles, CA), recorded using an acquisition board (OpenEphys, www.open-ephys.org), and saved on a personal computer at a sample rate of 30 kHz for offline spike sorting and analysis. After an experimental session was completed, the probe was moved approx. 65 µm along the middle cortical layers using the microdrive attached to the implant and the tissue was allowed to settle for at least 3 hours. Data collection could last up to four months, recording from 2-17 positions in each animal. Data collection was stopped when no primary-like responses could be detected anymore; criteria for primary-like responses were latency below 20 ms and reliable, clearly tuned responses to pure tone stimuli.

### Experimental setup

All experiments took place in a customized double-walled sound-attenuated chamber (workshop of the University of Oldenburg). Animals were monitored throughout experiments using an infrared camera (Pi NoIR, Raspberry Pi foundation, Cambridge, UK). Mice were placed on a custom horizontal and acoustically transparent running wheel made out of wire mesh and were free to run during the experiments. A loudspeaker (XT 300 K/4, Vifa, Viborg, Denmark) was attached 45 cm above the running wheel and delivered amplified sound (A-S501, Yamaha, Hamamatsu, Japan). Sound stimulation was generated digitally at a sample rate of 192 kHz using custom software written in MATLAB (MathWorks, Natick, MA, US) and were D/A converted by a USB sound device (Fireface UC, RME, Haimhausen, Germany). Both sound delivery and optogenetic manipulation were controlled using MATLAB. For activation of PV+ cells a blue laser was used (MDL-III-447 PSU-III-FDA, Changchun New Industries, Changchun, China), whereas deactivation of PV+ cells was done with an orange laser (Cobolt Mambo 50 594 nm, Cobolt AB, Solna, Sweden) using a shutter driver (Model SR474, Stanford Research Systems, Sunnyvale, CA, US). The optic fibres from both lasers were merged (TM105R3F1A, Thorlabs, Newton, NJ) and plugged onto the single optic fibre incorporated into the animal’s implant. After a block of auditory stimuli was presented, laser pulses activated or deactivated SSFO for the next stimulus block (see Fig. 1 A). The number of blocks for each SSFO condition varied depending on the stimulus protocol.

### Auditory stimuli

We used three auditory stimuli: (1) simple single pure tone stimuli (ST), (2) natural stimuli (NS), and (3) dynamic random chords (DRC). ST were presented randomly at 70 dB SPL between 4 and 70 kHz using eight frequencies per octave. The tones had a duration of either 0.5 or 1 s including 2 ms ramps and were generated with an inter-stimulus interval of 1 s or 2 s respectively. Each tone frequency was repeated three times within one stimulus block. The stimulus block was repeated ten times for each SSFO condition, resulting in at least 30 repetitions of each tone pulse. For NS, we extracted five natural animal vocalisations from an audio disc (‘Die Stimmen der Tiere 1 - Europa’, Cord Riechelmann, 2007; tracks: 13-tundra vole, 18-european pine vole, 40-yellowhammer, 45-european robin, 71-long eared owl) with a frequency spectrum centered between 4 and 25 kHz and up-sampled their sample rate to 192 kHz. The vocalisations were repeated in total 30 times in 3 to 6 blocks of each SSFO condition. DRC stimuli were similar to those described in previous studies (deCharms et al. 1998; Linden et al. 2003). The chords consisted of short tone pulses 20 ms in duration (including 5 ms ramps) with centre frequencies varying between 4.1 and 62 kHz in 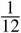-octave steps and sound levels varying between 25 and 70 dB SPL in 5 dB steps. Tone frequencies and levels were chosen randomly for each chord, with an average tone density of 2 tone pulses per octave. The DRC protocol lasted for 60 s and was repeated five times within a stimulus block. The stimulus block was repeated four times for both SSFO conditions.

## Data analysis

All analyses were performed offline using MATLAB, unless otherwise noted.

Continuous raw voltage traces were spike-detected and -sorted using a latent-variable spike-sorting algorithm (Sahani 1999) as previously described (Hildebrandt et al. 2017). For spike characterization, we set the waveform’s baseline 0.23 ms before peak (Keller et al. 2018) and normalized the unfiltered waveform with respect to the peak. We extracted the trough amplitude and measured spike width at 50% peak amplitude, tested for significant differences in medians between the subgroups using the Kruskal-Wallis test, and analyzed results further using the post-hoc Tukey-Kramer test. In order to compare optogenetic effect to the spike width, we used the Mann-Whitney U-Test.

### Criteria for exclusion and classification of units

For all of our analysis we only included units with a significant response during tone presentation. A unit with a significant response was defined as a unit for which the firing rate during tone presentation was significantly greater than spontaneous firing rate for at least three tested tone frequencies. Significance was tested by Student’s paired t-test to compare tone response with pre-tone firing on each of the repeated trials for each tone frequency in windows of 50ms (p<0.05 with Bonferroni correction for multiple tests). To classify the effect of PV+ activation on the neuronal responses, the mean firing rate over single sweeps including presentation of all tested frequencies was calculated, including tones and silent periods in between. Then, the distributions of the mean rate during single sweeps for both conditions were compared for each unit (Student’s paired t-test, single tail for decrease and enhancement respectively, p<0.05, with Bonferroni correction).

### Single tones (ST)

Onset responses for each tone frequency were computed as the mean response in a 40 ms window starting 15 ms after stimulus onset. The resulting tuning curves were smoothed using a Hanning window (width 3). For extracting the tuning width and the best frequency (BF), the tuning curves were normalized to their peaks. The tuning width was measured at 50% of the amplitude.

For correlation of tuning curves between control and PV-activated conditions, the tuning curves were normalized to the unit’s peak in the control condition. Slope and y-intercept values were extracted by using major-axis regression (i.e., two-dimensional least-mean-squares linear fit). In order to avoid floor effects, rates within the tuning curve lower than 10% of the maximum rate were discarded from the fit. In order to analyze the suppression of the tuning curves after PV+ cell activation, we calculated the value of the t-statistic *t*_val_ with 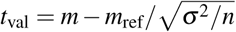, where *m* is the regression parameter (either slope or y-intercept), *m*_ref_ either 1 (slope) or 0 (y-intercept), *σ* the standard deviation of the regression parameter and *n* the number of points that went into the regression. The result was then compared to the t-distribution (single tail) and units categorized accordingly: divisive suppression in case of *p*_slope_ < 0.05 and *p*_intercept_ > 0.05; subtractive suppression for *p*_slope_ > 0.05 and *p*_intercept_ < 0.05.

### Dynamic random chords (DRC)

For each cell and SSFO condition, the average response to the DRC stimulus was used to estimate the respective spectro-temporal receptive fields (STRFs). STRFs were estimated with Automatic Smoothness Determination (Sahani and Linden 2003) within Python module ‘lnpy’ (https://github.com/arnefmeyer/lnpy). We set the dimensionality of the STRF to be 48 frequency channels and 15 time steps spanning 300 ms, chose a minimum STRF smoothness of 0.5 and tolerance of 10^−5^, and ran the optimisation for 100 iterations. For control trials, the spectral, temporal and overall smoothness scales were initialised at 4, 4 and 7 respectively. For PV-activated trials, the smoothness parameters were fixed at the optimised smoothness parameters obtained in the corresponding control trials, to ensure that comparisons between control and PV-activated trials were not confounded by differences in STRF smoothing parameters.

We included in the final DRC analysis all cells which were both: (1) responsive to pure tones (see above) and (2) responsive to the DRC stimulus (i.e., signal power of DRC response at least one standard error above zero (Sahani and Linden 2003; Williamson et al. 2016)).

We used the same major-axis regression procedure as with ST to compute correlations in STRF-predicted activity between control and PV-activated conditions.

## Recurrent network model and simulations

### Network model

We simulated an iso-frequency cortical column as a rate network model composed of *n*_*E*_ excitatory neurons and *n*_*I*_ inhibitory neurons with the following dynamics:

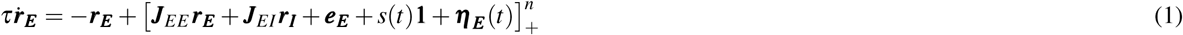

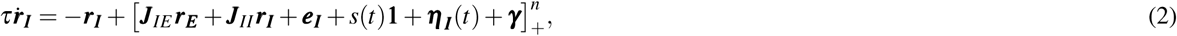

where *τ* is the effective rate time constant, ***J***_*AB*_ the connectivity matrix onto population *A* from population *B*, ***e***_***A***_ is the tonic background input different for every neuron in population *A, s*(*t*) is the phasic thalamic input and is the same for every neuron, ***η*** _***A***_(*t*) is the white noise background input different for every neuron in population *A*, ***γ*** is the optogenetic input into inhibitory neurons in PV-activated trials and is different for every neuron, and 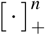 is a threshold-linear input-output function raised to the power *n* and operating elementwise.

The connectivity matrix ***J***_*AB*_ was tuned to have sparsity *ρ*:

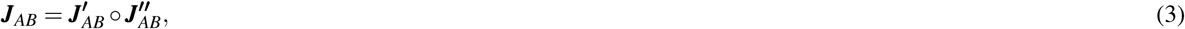

where ∘ denotes the Hadamard product between the weight matrix 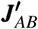 and the sparsity matrix 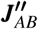. Within 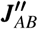, each element is sampled from a Bernoulli distribution Bern(1 − *ρ*).

### Thalamic input

In the model we assume all cortical neurons receive the same thalamic input *s*(*t*), without loss of generality. The input onset is at *t*_*on*_ and is active for a time window Δ*t*. The input tuning is bell-shaped with respect to sound frequency, with centre frequency *f*_*c*_ and standard deviation *σ*_*f*_, and *n*_*f*_ sound frequencies overall:

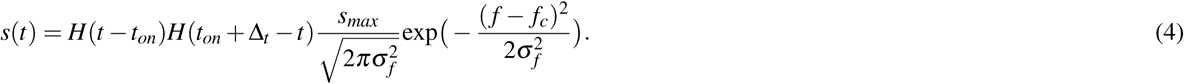

where *H*(·) is the Heaviside (step) function.

### Numerical simulations

The network model was simulated in MATLAB using the forward Euler-Maruyama for each of the *n*_*f*_ thalamic inputs, and for each of two conditions, control and PV-activated. The parameters used in the simulations are listed in Table 1.

**Table 1.**
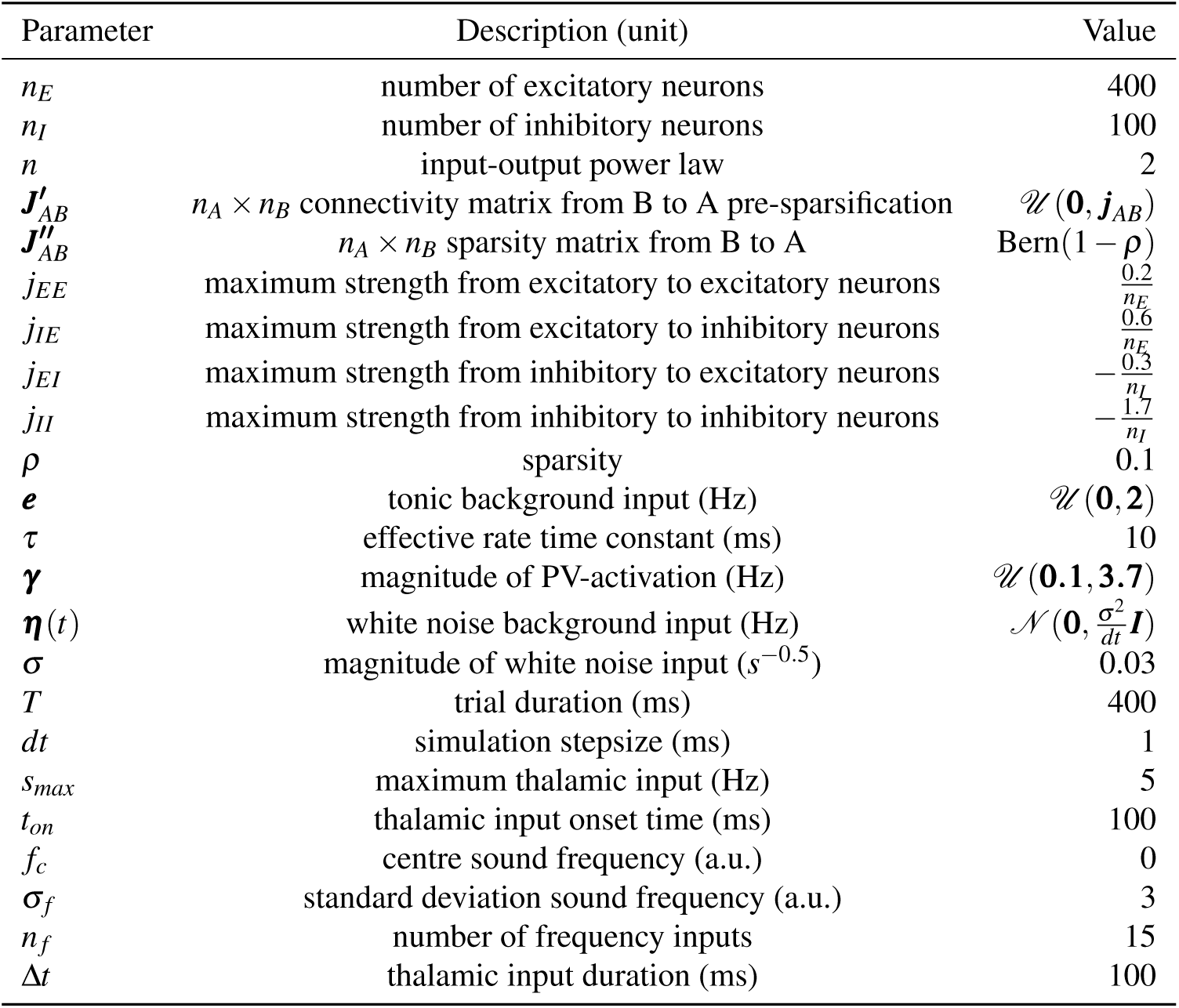
Simulation parameters.

The response to a tone was computed as the average response in a 90 ms window following the tone onset.

## Author contributions

Conceptualization: J.F.L, M.S. and K.J.H. Methodology: T.G. and P.J.G. Software: T.G., P.J.G. and K.J.H. Formal Analysis: T.G. and P.J.G. Investigation: T.G. and K.J.H. Writing – Original Draft: T.G., P.J.G. and K.J.H. Writing – Review Editing: T.G., P.J.G., J.F.L., M.S. and K.J.H. Supervision: J.F.L, M.S. and K.J.H. Funding Acquisition: J.F.L, M.S. and K.J.H.

## Acknowledgements

We thank H. Hosseini and D. Gonschorek for help with data collection; K. Deissoroth for advice on opsin selection and providing the constructs; C. Thiel for comments on an earlier version of the manuscript; and S. Meiser and M. Rogalla for general support and helpful discussions. Funding for this work was provided by the DFG Cluster of Excellence EXC 644 1077/1 “Hearing4all” (K.J.H.), the Biological Sciences and Biotechnology Research Council (BB/P7201/1, J.F.L.), the Gatsby Charitable Foundation (GAT3528; M.S.), and the Simons Foundation (SCGB323228 and SCGB543039; M.S.).

## Supplementary material

**Figure S1.**
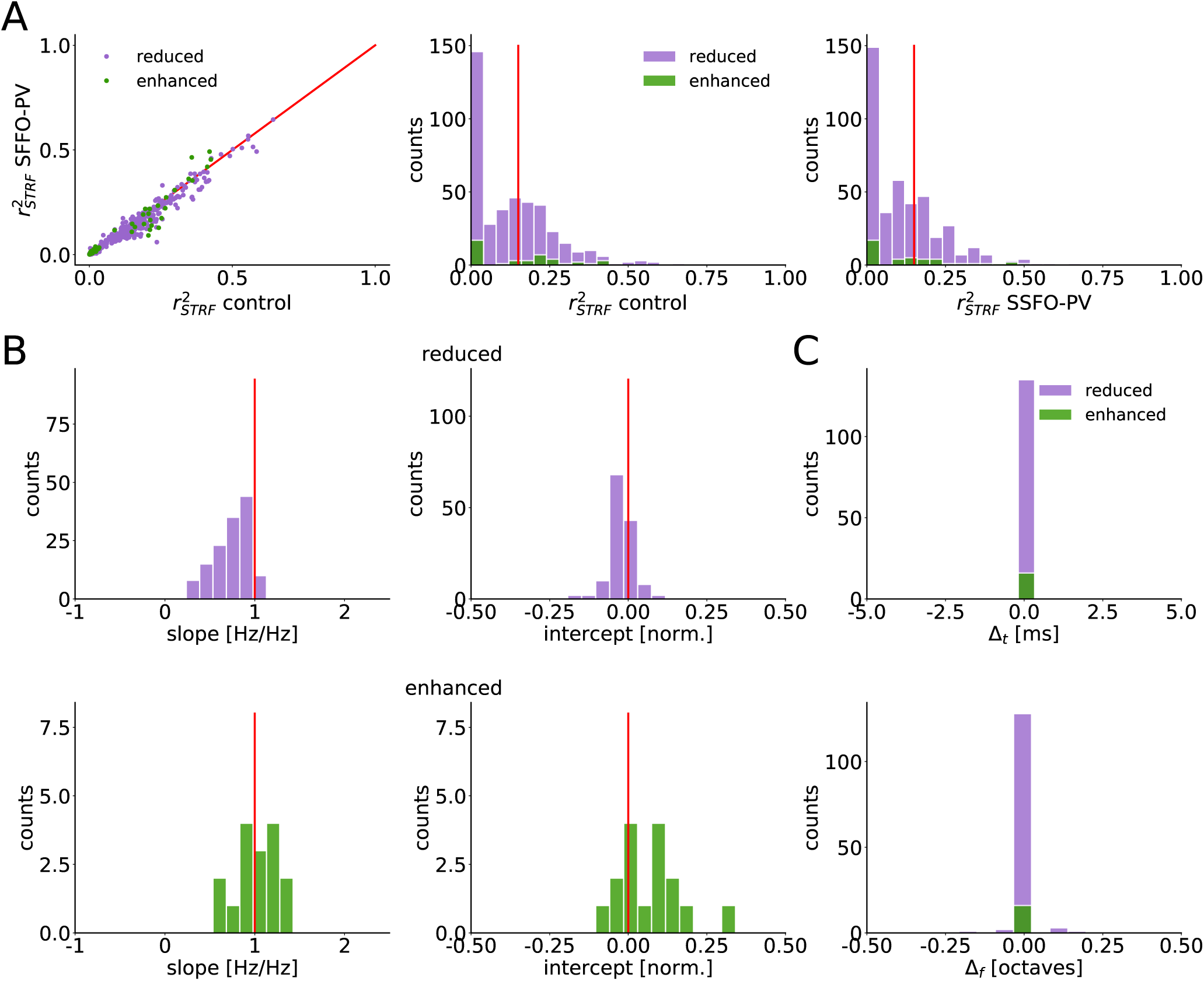
Units with coefficient of determination *r*^2^ > 0.15 in STRF estimation have STRFs which are divisively scaled by PV-activation, while preserving their structure. (**A**): Coefficients of determination of STRF estimation in PV-activated versus control conditions (left). Coefficients of determination are highly correlated (*ρ* ≈ 0.976) and have similar range in PV-activated versus control conditions. Note that 1% of the reduced units and 5% of the enhanced units have *r*^2^ < 0 in PV-activated condition and are excluded for better visualisation of the bulk of the data. (**B**): For reduced (top) and enhanced (bottom) units, slopes (left) and intercepts (right) of linear fits to STRF predicted activities in PV-activated versus control conditions. Only units with *r*^2^ > 0.15 are included (enhanced units n = 16 (out of a total 44) and reduced units n = 135 (out of a total 419)). Results are robust to the choice of *r*^2^ threshold (results not shown). (**C**): For most units, sustained PV-activation does not shift the STRF in time or frequency.

**Figure S2.**
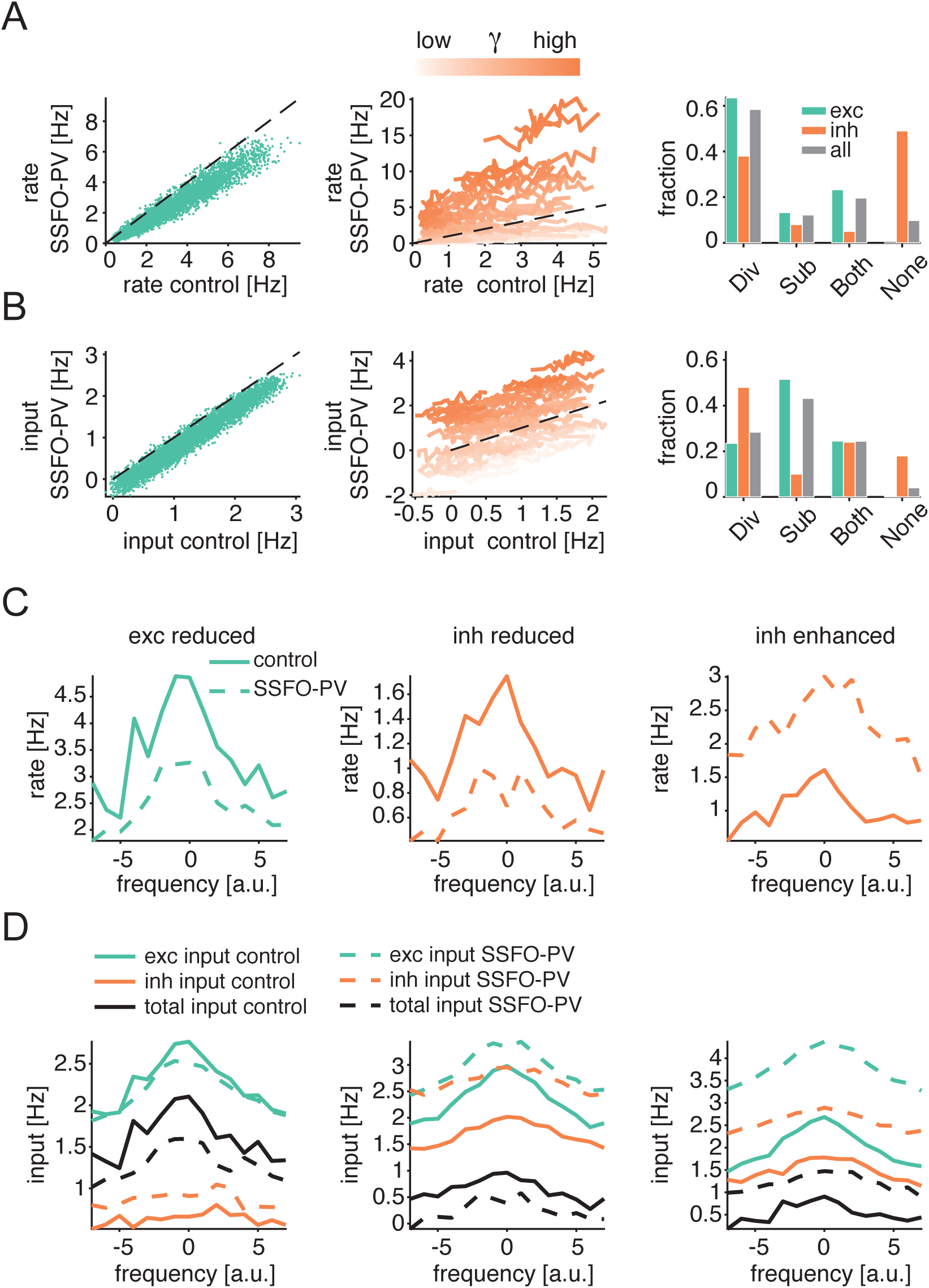
Network model: divisive modulation of firing rates and more subtractive modulation of respective inputs. (**A**): Model tuning curves of excitatory (left) and inhibitory (center) cells in control and PV-activated conditions. Excitatory neural activity is typically divisively modulated in PV-activated compared to control conditions (right). Inhibitory cell activity shows a broad range of effects, i.e. divisive, additive and multiplicative effects, depending on the magnitude of direct PV-activation. (**B**): Tuning of the inputs to the excitatory (left) and inhibitory (center) cells in control and PV-activated conditions. Inputs to excitatory neurons are more subtractively modulated compared to excitatory firing rates (right). (**C**): Three representative tuning curves in control and PV-activated conditions: excitatory (left) and inhibitory (middle) cells divisively reduced during PV-activation, and inhibitory cell (right) enhanced during PV-activation. (**D**) Tuning of the inputs for the same cells as in (C).

